# Use of multiple picosecond high-mass molecular dynamics simulations to predict crystallographic B-factors of folded globular proteins

**DOI:** 10.1101/052126

**Authors:** Yuan-Ping Pang

## Abstract

Predicting crystallographic B-factors of a protein from a conventional molecular dynamics simulation is challenging in part because the B-factors calculated through sampling the atomic positional fluctuations in a picosecond molecular dynamics simulation are unreliable and the sampling of a longer simulation yields overly large root mean square deviations between calculated and experimental B-factors. This article reports improved B-factor prediction achieved by sampling the atomic positional fluctuations in multiple picosecond molecular dynamics simulations that use uniformly increased atomic masses by 100-fold to increase time resolution. Using the third immunoglobulin-binding domain of protein G, bovine pancreatic trypsin inhibitor, ubiquitin, and lysozyme as model systems, the B-factor root mean square deviations (mean ± standard error) of these proteins were 3.1 ± 0.2–9 ± 1 Å^2^ for Cα and 7.3 ± 0.9–9.6 ± 0.2 Å^2^ for Cγ, when the sampling was done, for each of these proteins, over 20 distinct, independent, and 50-picosecond high-mass molecular dynamics simulations using AMBER forcefield FF_12_MC or FF_14_SB. These results suggest that sampling the atomic positional fluctuations in multiple picosecond high-mass molecular dynamics simulations may be conducive to *a priori* prediction of crystallographic B-factors of a folded globular protein.

## 1. Introduction

The B-factor (also known as the Debye-Waller factor or B-value) of a given atom in a crystal structure is defined as 8π^2^〈*u*^2^〉 that is used in refining the crystal structure to reflect the displacement *u* of the atom from its mean position in the crystal structure (*viz.*, the uncertainty of the atomic mean position) [1-10]. The displacement *u* attenuates X-ray scattering and is caused by the thermal motion, conformational disorder, and static lattice disorder of the atom [6]. It is worth noting that the experimentally determined B-factor is not a quantity that is directly observed from an experiment. Instead, it is a function that not only decreases as the resolution of the crystal structure increases [10] but also depends on the restraints that are applied on B-factors in refining the crystal structure [4,8]. B-factors can be unrealistic if excessive refinement is performed to achieve a higher resolution. B-factors of one crystal structure cannot be compared to those of another without detailed knowledge of the refinement processes for the two comparing structures. It is also worthy of noting that the Subcommittee on Atomic Displacement Parameter Nomenclature recommends avoiding referring to B-factor as “temperature factor” in part because the displacement may not be caused entirely by the thermal motion [7].

Despite the complex nature of B-factor and challenges of separating the thermal motion in time from the conformational and static lattice disorders in space [11], B-factors of a protein crystal structure can be used to quantitatively identify *less mobile* regions of a crystal structure as long as the structure is determined without substantial crystal lattice defects, rigid-body motions, and refinement errors [8,12,13]. A low B-factor indicates low thermal motion, while a high B-factor may imply high thermal motion. Normalized main-chain B-factors of a protein have been used as an estimator of flexibility for each residue of the protein [14-19] to offer useful information for drug-target identification. Unscaled main-chain and side-chain B-factors of a protein can be used to identify ordered regions of a folded globular protein and relatively rigid side chains of active-site residues for target-structure–based drug design [20,21]. Other uses of B-factors are outlined in Ref. [22].

As of August 2016, there are more than 65 million protein sequences at the Universal Protein Resource (http://www.uniprot.org/statistics/TrEMBL) compared to about 106 thousand protein crystal structures available at the Protein Data Bank (http://www.rcsb.org/pdb/statistics/holdings.do). This difference indicates that one can use crystallographic methods to determine structures and B-factors of only a fraction of known protein sequences. Most known protein sequences will have to be used for target identification and drug design through generation and refinement of comparative or homology models from the protein sequences [23-42]. Currently, knowledge-based methods can predict main-chain B-factor distribution of a protein from either its sequence using statistical methods [15,17-19,43-46] or from its structure using a single-parameter harmonic potential [47,48] with Pearson correlation coefficients up to 0.71 for the predicted B-factors relative to the experimental values. These methods do not require intense computation and can rapidly predict B-factors of large numbers of protein sequences to facilitate the use of these sequences in drug-target identification. However, target-structure–based drug design requires more detailed B-factor information than drug-target identification. To design drug candidates whose binding to their protein targets is both enthalpy- and entropy-driven, one needs the information on side-chain motions of active-site residues in a protein target. Prediction of side-chain B-factors by the knowledge-based methods has not been reported to date and may not be feasible through the use of a single-parameter harmonic potential that is inapplicable to high frequency modes pertaining to rapid oscillations of some amino acid side chains [49].

To complement the current knowledge-based methods, there is a need to develop physics-based methods for predicting unscaled B-factors of both main-chain and side-chain atoms of a protein crystal structure or a refined comparative protein model from molecular dynamics (MD) simulation. By solving the Newtonian equations of motion for all atoms in a molecular system as a function of time, MD simulation is a general method to simulate atomic motions of the system for insights into dynamical properties of the system such as transport coefficients, time-dependent response to perturbations, rheological properties, and spectra [50]. However, predicting B-factors of a folded globular protein by sampling the atomic positional fluctuations of a protein in a conventional MD simulation with solvation may not be feasible because of the use of different protein environments, different timescales to detect thermal motions, and different methods to determine B-factors [51]. For example, a reported MD simulation study showed that the B-factors derived on the picosecond timescale were unreliable and that the simulated B-factors on the nanosecond timescale were considerably larger than the experimental values [51]. Although simulations of proteins in their crystalline state [52,53] can avoid the difference in protein environment, such simulations are inapplicable to *a priori* B-factor prediction.

This article reports an evaluation study of a physics-based method that samples the atomic positional fluctuations in 20 distinct, independent, unrestricted, unbiased, picosecond, and classical isobaric–isothermal (NPT) MD simulations with uniformly scaled atomic masses to predict *a priori* main-chain and side-chain B-factors of a folded globular protein for target-structure–based drug design. The model systems of folded globular proteins used in this study were the third immunoglobulin-binding domain of protein G (GB3; PDB ID: 1IGD; resolution: 1.10 Å) [54], bovine pancreatic trypsin inhibitor (BPTI; PDB ID: 4PTI; resolution: 1.50 Å) [55], ubiquitin (PDB ID: 1UBQ; resolution: 1.80 Å) [56], and lysozyme (PDB ID: 4LZT; resolution: 0.95 Å) [57]. Two distinct AMBER forcefields, FF_12_MC [42,58-60] and FF_14_SB [61], were used to evaluate the method in a forcefield-independent manner. The root mean square deviations (RMSDs) and Pearson correlation coefficients (PCCs) between the experimental B-factors and the predicted values by the physics-based method were compared respectively to the estimated standard error of the experimental B-factors derived from the refinement procedure [8] and to the PCCs of the reported knowledge-based methods [46,47] in order to assess the quality of the B-factors predicted by the physics-based method. Unless otherwise specified below, all B-factors are unscaled, and all simulations are multiple, distinct, independent, unrestricted, unbiased, and classical NPT MD simulations.

## 2. Theory

### 2.1. Using uniformly reduced atomic masses to compress the MD simulation time

Reducing atomic masses of the entire simulation system (including both solute and solvent) uniformly by tenfold—hereafter referred to as low masses—can enhance configurational sampling in NPT MD simulations [62]. The effectiveness of the low-mass NPT MD simulation technique can be explained as follows: To determine the relative configurational sampling efficiencies of two simulations of the same molecule—one with standard masses and another with low masses, the units of distance [*l*] and energy [*m*]([*l*]/[*t*])^2^ of the low-mass simulation are kept identical to those of the standard-mass simulation, noting that energy and temperature have the same unit. This is so that the structure and energy of the low-mass simulation system can be compared to those of the standard-mass simulation system. Let superscripts ^lmt^and ^smt^denote the times for the low-mass and standard-mass systems, respectively. Then [*m*^lmt^] = 0.1 [*m*^smt^], [*l*^lmt^] = [*l*^smt^], and [*m*^lmt^]([*l*^lmt^]/[*t*^lmt^])^2^ = [*m*^smt^]([*l*^smt^]/[*t*^smt^])^2^ lead to 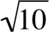 [*t*^lmt^] = [*t*^smt^]. A conventional MD simulation program takes the timestep size (Δt) of the standard-mass time rather than that of the low-mass time. Therefore, low-mass MD simulations at Δt = 1.00 fs^smt^ (viz., 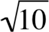 fs^lmt^) are theoretically equivalent to standard-mass MD simulations at 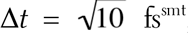, as long as both standard-mass and low-mass simulations are carried out for the same number of timesteps and there are no precision issues in performing these simulations. This equivalence of mass downscaling and timestep-size upscaling explains why uniform mass reduction can compress the MD simulation time and why low-mass NPT MD simulations at Δt = 1.00 fs^smt^ can offer better configurational sampling efficacy than conventional standard-mass NPT MD simulations at *Δt* = 1.00 fs^smt^ or *Δt* = 2.00 fs^smt^. It also clarifies why the kinetics of the low-mass simulation system can be converted to the kinetics of the standard-mass simulation system simply by scaling the low-mass time with a factor of 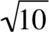 [60]. Further, this equivalence explains there are limitations on the use of the mass reduction technique to improve configurational sampling efficiency. Lengthening the timestep size inevitably reduces integration accuracy of an MD simulation. However, the integration accuracy reduction caused by a timestep-size increase is temperature dependent. Therefore, to avoid serious integration errors, low-mass NPT MD simulations must be performed with the double-precision floating-point format and at *Δt* ≤1.00 fs^smt^ and a temperature of ≤340 K [60]. Because temperatures of biological systems rarely exceed 340 K and because MD simulations are performed typically with the double-precision floating-point format, low-mass NPT MD simulation is a viable configurational sampling enhancement technique for protein simulations at a temperature of ≤340 K. In this context, to efficiently sample alternative conformations from a crystallographically determined conformation, low-mass NPT MD simulations at *Δt* = 1.00 fs^smt^ and temperature of <340 K were used for GB3, BPTI, ubiquitin, and lysozyme in this study.

### 2.2. Using uniformly increased atomic masses to expand the MD simulation time

In the same vein, let superscript ^hmt^ denote the time for the system with uniformly increased atomic masses by 100-fold (hereafter referred to as high masses), then [*m*^hmt^] = 100 [*m*^smt^], [*l*^hmt^] = [*l*^smt^], and [*m*^hmt^]([*l*^hmt^]/[*t*^hmt^])^2^ = [*m*^smt^]([*l*^smt^]/[*t*^smt^])^2^ lead to [t^hmt^] = 10 [t^smt^]. This equivalence of mass upscaling and timestep-size downscaling explains why uniform mass increase can expand the MD simulation time and why high-mass NPT MD simulations at *Δt* = 1.00 fs^smt^ can increase their time resolution by tenfold. Therefore, to adequately sample the atomic positional fluctuations in a short simulation, high-mass NPT MD simulations at *Δt* = 1.00 fs^smt^ were used for GB3, BPTI, ubiquitin, and lysozyme in the present study. Although standard-mass simulations at *Δt* = 0.10 fs^smt^ can achieve the same time resolution, the high-mass simulation with *Δt* = 1.00 fs^smt^ has an advantage in that, through modifying the atomic masses specified in a forcefield parameter file rather than the source code of the simulation package, one can simulate a guest**•**host complex with the compressed and expanded simulation times respectively applied to the guest and the host or a homology model of a protein with the compressed and expanded simulation times respectively applied to the active-site region and the rest of the protein. The simulation time resolution can also be increased by sampling conformations saved at every 50 timesteps of a standard-mass simulation at *Δt* = 1.00 fs^smt^ [4,51] rather than sampling conformations saved at every 10^3^ timesteps of a high-mass simulation as described in Section 3.2. However, to simultaneously perform 20 simulations of a large protein with explicit solvation, the high-mass simulations are preferred over the standard-mass simulations because simultaneously saving 20 large files of the coordinates of the protein with a vast number of water molecules at every 50 timesteps is more computationally expensive than at every 10^3^ timesteps.

## 3. Methods

### 3.1. MD simulations of folded globular proteins

A folded globular protein was solvated with the TIP3P water [63] with surrounding counter ions and then energy-minimized for 100 cycles of steepest-descent minimization followed by 900 cycles of conjugate-gradient minimization to remove close van der Waals contacts using SANDER of AMBER _11_ (University of California, San Francisco). The resulting system was heated—in 20 distinct, independent, unrestricted, unbiased, and classical MD simulations with a periodic boundary condition and unique seed numbers for initial velocities—from 0 to 295 or 297 K at a rate of 10 K/ps under constant temperature and constant volume, then equilibrated with a periodic boundary condition for 10^6^ timesteps under constant temperature and constant pressure of 1 atm employing isotropic molecule-based scaling, and lastly simulated under the NPT condition at 1 atm and a constant temperature of 295 K or 297 K using PMEMD of AMBER 11.

The initial conformations of GB_3_, BPTI, ubiquitin, and lysozyme for the simulations were taken from the crystal structures of PDB IDs of _1_IGD, _5_PTI, _1_UBQ, and _4_LZT, respectively. A truncated _1_IGD structure (residues 6–61) was used for the GB_3_ simulations. Four interior water molecules (WAT_111_, WAT_112_, WAT_113_, and WAT_122_) were included in the initial 5PTI conformation. The simulations for GB_3_, BPTI, and ubiquitin were done at _297_ K as the exact data-collection temperatures of these proteins had not been reported. The lysozyme simulations were done at the reported data-collection temperature of _295_ K [57].

The numbers of TIP_3_P waters and surrounding ions, initial solvation box size, and protonation states of ionizable residues used for the NPT MD simulations are provided in Table 1. The 20 unique seed numbers for initial velocities of Simulations 1–20 were taken from Ref. [58]. All simulations used (*i*) a dielectric constant of 1.0, (*ii*) the Berendsen coupling algorithm [64], (*iii*) the Particle Mesh Ewald method to calculate electrostatic interactions of two atoms at a separation of >8 Å [65], (*iv*) *Δt* = 1.00 fs^smt^, (*v*) the SHAKE-bond-length constraints applied to all bonds involving hydrogen, (*vi*) a protocol to save the image closest to the middle of the “primary box” to the restart and trajectory files, (*vii*) a formatted restart file, (*viii*) the revised alkali and halide ions parameters [66], (*ix*) a cutoff of 8.0 Å for nonbonded interactions, (*x*) the atomic masses of the entire simulation system (including both solute and solvent) that were uniformly increased by 100-fold or decreased by tenfold relative to the standard atomic masses, and (*xi*) default values of all other inputs of the PMEMD module. The forcefield parameters of FF_12_MC are available in the Supporting Information of Ref. [60]. All simulations were performed on a cluster of 100 12-core Apple Mac Pros with Intel Westmere (2.40/2.93 GHz).

### 3.2. Crystallographic B-factor prediction

Using a two-step procedure with PTRAJ of AmberTools 1.5, the B-factors of Cα and Cγ atoms in a folded globular protein were predicted from all conformations saved at every 10^3^ timesteps of 20 simulations of the protein using the simulation conditions described above. The first step was to align all saved conformations onto the first saved one to obtain an average conformation using root mean square fit of all CA atoms (for Cα B-factors) or all CG and CG2 atoms (for Cγ B-factors). The second step was to root mean square fit all CA atoms (or all CG and CG2 atoms) in all saved conformations onto the corresponding atoms of the average conformation and then calculate the Cα (or Cγ) B-factors using the “atomicfluct” command in PTRAJ. For each protein, the calculated B-factor of an atom in Fig. 1 and Table S1 of Supplementary Material was the mean of all B-factors of the atom derived from 20 simulations of the protein. The standard error (SE) of a B-factor was calculated according to Eq. 2 of Ref. [59]. The SE of an RMSD between computed and experimental B-factors was calculated using the same method for the SE of a B-factor. The experimental B-factors of GB3, BPTI, ubiquitin, and lysozyme were taken from the crystal structures of PDB IDs of 1IGD, 4PTI, 1UBQ, and 4LZT, respectively.

**Fig. 1.**
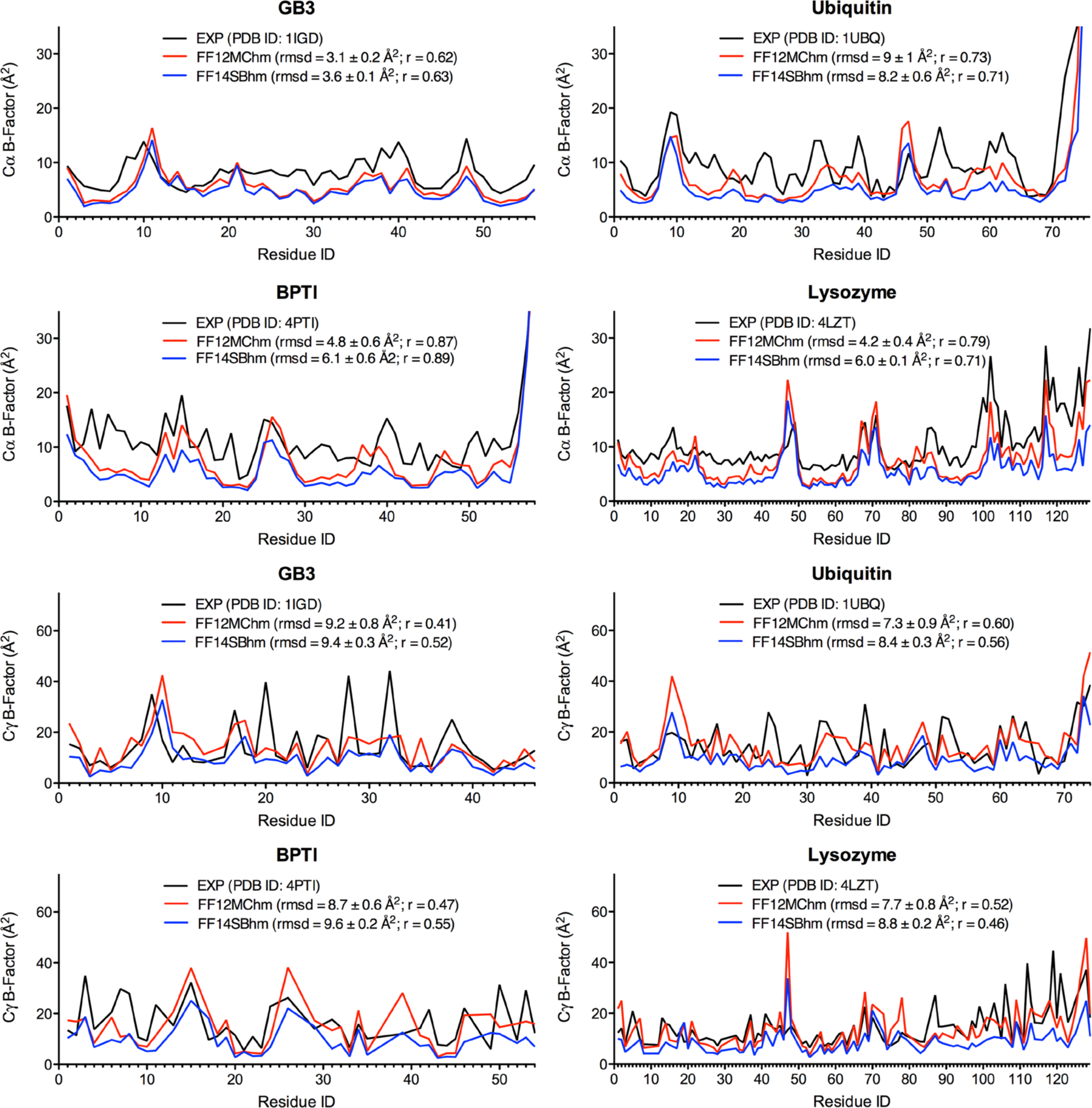
Experimental and calculated B-factors of GB3, BPTI, ubiquitin, and lysozyme. The B-factors were calculated from 20 50-ps^smt^ high-mass molecular dynamics simulations using FF_12_MChm or FF_14_SBhm. The letter “r” is the abbreviation for the Pearson correlation coefficient.

### 3.3. Correlation analysis

PCCs were obtained from correlation analysis using PRISM 5 for Mac OS X of GraphPad Software (La Jolla, California) with the assumption that data were sampled from Gaussian populations.

## 4. Results and discussion

### 4.1. Using high–time-resolution picosecond MD simulations to calculate B-factors

The internal motions—such as the motions of backbone N–H bonds of a folded globular protein at the solution state—are on the order of tens or hundreds of ps^smt^ [67]. Therefore, the timescale of the thermal motions reflected in the B-factors of a protein at the crystalline state is unlikely greater than a nanosecond. As described in Section 1, the B-factor of a given atom reflects both the thermal motion and the conformation and static lattice disorders of the atom [6]. In this context, 20 high-mass MD simulations of a folded globular protein were carried out to investigate whether combining the sampling of the atomic positional fluctuations of the protein on a timescale of tens or hundreds of ps^smt^ with the sampling of such fluctuations over conformations derived from the 20 simulations could approximate the experimental B-factors. High-mass simulations were used to increase the time resolution of the simulations and performed with FF_12_MChm or FF_14_SBhm, which denote the AMBER forcefields FF_12_MC or FF_14_SB with all atomic masses that were uniformly increased by 100-fold relative to the standard atomic masses.

As listed in Table 2, regardless of which forcefield was used, the RMSDs between computed and experimental B-factors of Cα and Cγ were <10 Å^2^ for all four proteins when the atomic positional fluctuations of these proteins were sampled on the timescale of 50 ps^smt^. When FF_12_MChm was used, longer samplings led to the RMSDs of ≥10 Å^2^ for all four proteins, and these RMSDs progressed in time (Table 2). When FF_14_SBhm was used with longer samplings, the RMSDs were also >10 Å^2^ for GB3, ubiquitin, and BPTI. For the lysozyme B-factors predicted with FF_14_SBhm, the RMSDs were <10 Å^2^ when the sampling were done on the timescale of <1 ns^smt^, but the RMSDs were >15 Å^2^ for the samplings on the timescale of 10 or 20 ns^smt^ (Table 2). FF_12_MChm best reproduced most of the experimental B-factors on the timescale of 50 ps^smt^ with RMSDs (mean ± SE) ranging from 3.1 ± 0.2 to 9 ± 1 Å^2^ for Cα and from 7.3 ± 0.9 to 9.2 ± 0.8 Å^2^ for Cγ. FF_14_SBhm also best reproduced most of the experimental B-factors on the timescale of 50 ps^smt^ with RMSDs (mean ± SE) from 3.6 ± 0.1 to 8.2 ± 0.6 Å^2^ for Cα and from 8.4 ± 0.3 to 9.6 ± 0.2 Å^2^ for Cγ. Regardless of which forcefield was used, the means and SEs of the B-factor RMSDs of ubiquitin were larger than those of the other proteins (Table 2). It was logical to suspect that the conformational variations resulting from 20 simulations might be insufficient to adequately represent the conformational disorder of the ubiquitin crystals. However, increasing the number of the ubiquitin simulations from 20 to 40 or 80 only reduced the SEs but not the means (Table 3).

**Table 1.** Numbers of TIP3P waters and ions, initial solvation box size, and protonation state of ionizable residue used in molecular dynamics simulations.

| Sequence | # of H<sub>2</sub>O | # of Na<sup>+</sup> | # of Cl<sup>-</sup> | Box size (Å<sup>3</sup>) | Expt pH | Protonation State of Ionizable Residue |
| --- | --- | --- | --- | --- | --- | --- |
| GB3 | 2528 | 2 | 0 | 45x57x47 | 5.8 | ASP, GLU, LYS |
| BPTI | 3108 | 0 | 6 | 47x46x65 | 4.6 | ARG, ASP, GLU, LYS |
| Ubiquitin | 3881 | 0 | 1 | 50x66x53 | 4.7 | ARG, ASP, GLU, LYS, HIP |
| Lysozyme | 5849 | 0 | 12 | 60x61x69 | 3.8 | ARG, ASP, ASH<sup>101</sup>, GLH, LYS, HIP |

**Table 2.** Root mean square deviations between experimental and calculated B-factors of GB3, BPTI, ubiquitin, and lysozyme.

| Protein<br>(temperature) | Time<br>(ps<sup>smt</sup>) | RMSD (mean ± SE in $\text{\AA}^2$) | | | |
| --- | --- | --- | --- | --- | --- |
| | | Cα | | Cγ | |
|  |  | FF12MChm | FF14SBhm | FF12MChm | FF14SBhm |
| Ubiquitin<br>(297 K) | 25 | 6.2±0.3 | 7.1±0.2 | 7.0±0.9 | 9.3±0.2 |
| | 50 | 9±1 | 8.2±0.6 | 7.3±0.9 | 8.4±0.3 |
| | 100 | 16±2 | 12±1 | 12±1 | 7.8±0.6 |
| | 200 | 32±3 | 21±2 | 20±2 | 9±1 |
| | 300 | 37±4 | 28±3 | 25±3 | 10±1 |
| | 400 | 40±4 | 32±3 | 27±3 | 11±1 |
| | 500 | 43±4 | 36±3 | 29±3 | 12±2 |
| BPTI<br>(297 K) | 25 | 5.9±0.3 | 6.8±0.3 | 8.6±0.4 | 10.7±0.2 |
| | 50 | 4.8±0.6 | 6.1±0.6 | 8.7±0.6 | 9.6±0.2 |
| | 100 | 5.2±0.8 | 7.3±0.9 | 11±1 | 9.1±0.3 |
| | 200 | 8±1 | 10±1 | 13±1 | 8.8±0.4 |
| | 300 | 13±2 | 14±2 | 15±1 | 8.8±0.5 |
| | 400 | 15±2 | 16±2 | 17±1 | 8.9±0.6 |
| | 500 | 17±2 | 18±2 | 19±1 | 8.9±0.6 |
| GB3<br>(297 K) | 25 | 3.7±0.1 | 4.2±0.1 | 9.3±0.5 | 10.3±0.2 |
| | 50 | 3.1±0.2 | 3.6±0.1 | 9.2±0.8 | 9.4±0.3 |
| | 100 | 3.7±0.7 | 3.4±0.2 | 12±2 | 8.8±0.6 |
| | 200 | 5.3±0.9 | 3.3±0.2 | 17±2 | 8.4±0.7 |
| | 300 | 5.9±0.8 | 3.2±0.2 | 19±2 | 8.0±0.6 |
| | 400 | 8±1 | 3.3±0.2 | 23±2 | 8.4±0.6 |
| | 500 | 9±1 | 3.6±0.3 | 25±2 | 9.5±0.9 |
| | 600 | 9±1 | 4.0±0.5 | 26±2 | 11±1 |
| | 700 | 10±1 | 4.3±0.5 | 27±2 | 12±1 |
| | 800 | 10±1 | 4.6±0.6 | 28±2 | 12±1 |
| | 900 | 10±1 | 4.9±0.7 | 28±2 | 13±2 |
| | 1000 | 10±1 | 5.2±0.7 | 29±2 | 13±2 |
| Lysozyme<br>(295K) | 25 | 5.2±0.3 | 6.7±0.1 | 7.4±0.5 | 9.5±0.1 |
| | 50 | 4.2±0.4 | 6.0±0.1 | 7.7±0.7 | 8.8±0.2 |
| | 100 | 3.5±0.6 | 5.5±0.1 | 10±1 | 8.4±0.2 |
| | 200 | 4.0±0.6 | 5.1±0.1 | 13±1 | 8.3±0.3 |
| | 300 | 5.2±0.6 | 5.1±0.1 | 17±1 | 8.6±0.4 |
| | 400 | 6.9±0.8 | 5.0±0.1 | 20±1 | 8.9±0.4 |
| | 500 | 8±1 | 4.9±0.1 | 22±2 | 9.0±0.4 |
| | 600 | 9±1 | 4.9±0.1 | 24±2 | 9.2±0.4 |
| | 700 | 10±1 | 4.9±0.1 | 26±3 | 9.4±0.4 |
| | 800 | 11±2 | 4.8±0.1 | 27±3 | 9.5±0.4 |
| | 900 | 11±2 | 4.8±0.1 | 28±3 | 9.6±0.4 |
| | 1000 | 12±2 | 4.8±0.1 | 29±3 | 9.7±0.4 |
| | 10,000 | — | 4.7±0.5 | — | 16±1 |
| | 20,000 | — | 5.4±0.8 | — | 19±2 |

Time: the duration of 20 distinct, independent, unrestricted, unbiased, and isobaric–isothermal molecular dynamics simulations over which the B-factors were calculated. RMSD: root mean square deviation. SE: standard error calculated from 20 distinct, independent, unrestricted, unbiased, and isobaric–isothermal molecular dynamics simulations.

**Table 3.** Effects of the number of molecular dynamics simulations on the root mean square deviation between experimental and calculated B-factors of ubiquitin.

| Forcefield | Time<br>(ps<sup>smt</sup>) | RMSD (mean ± SE in $\text{\AA}^2$) | | | | | |
| --- | --- | --- | --- | --- | --- | --- | --- |
| | | Cα | | | Cγ | | |
|  |  | N = 20 | N = 40 | N = 80 | N = 20 | N = 40 | N = 80 |
| FF12MChm | 25 | 6.2±0.3 | 6.5±0.3 | 6.7±0.2 | 7.0±0.9 | 7.0±0.5 | 7.2±0.3 |
| | 50 | 9±1 | 9.2±0.8 | 9.4±0.6 | 7.3±0.9 | 7.4±0.6 | 7.2±0.4 |
| | 100 | 16±2 | 15±1 | 15.0±0.8 | 12±1 | 11.0±0.8 | 10.5±0.6 |
| FF14SBhm | 25 | 7.1±0.2 | 7.3±0.3 | 7.4±0.3 | 9.3±0.2 | 9.4±0.1 | 9.3±0.1 |
| | 50 | 8.2±0.6 | 8.3±0.7 | 8.2±0.5 | 8.4±0.3 | 8.3±0.2 | 8.2±0.1 |
| | 100 | 12±1 | 11.0±0.9 | 13±1 | 7.8±0.6 | 7.6±0.4 | 7.5±0.3 |

Time: the duration of N distinct, independent, unrestricted, unbiased, and isobaric–isothermal molecular dynamics simulations over which the B-factors were calculated. RMSD: root mean square deviation. SE: standard error calculated from N number of distinct, independent, unrestricted, unbiased, and isobaric–isothermal molecular dynamics simulations.

For all four proteins, the agreement of the calculated Cα and Cγ B-factors on the timescale of 50 ps^smt^ with the experimental values is shown in Fig. 1, and the SEs of the predicted B-factors shown in Fig. 1 are listed in Table S1 of Supplementary Material. The B-factor RMSDs (mean ± SE) of these proteins using both FF_12_MChm and FF_14_SBhm ranged from 3.1 ± 0.2 to 9 ± 1 Å^2^ for Cα and from 7.3 ± 0.9 to 9.6 ± 0.2 Å^2^ for Cγ (Fig. 1). The respective PCCs were 0.62–0.87 or 0.63–0.89 for the Cα B-factors of the four proteins that were predicted using FF_12_MChm or FF_14_SBhm relative to the experimental B-factors (Fig. 1). The PCCs of the predicted Cγ B-factors using FF_12_MChm or FF_14_SBhm were 0.41–0.60 or 0.46–0.56 for the four proteins, respectively (Fig. 1). The average PCCs of the predicted B-factors using FF_12_MC and FF_14_SB were 0.75 and 0.74 for Cα and 0.50 and 0.52 for Cγ, respectively. These results suggest that combining the sampling of the atomic positional fluctuations over the ~50-ps^smt^ timescale with the sampling of such fluctuations over conformations derived from 20 distinct ~50-ps^smt^ simulations can approximate the experimental B-factors with RMSDs of <10 Å^2^ and the PCCs of 0.62–0.89 for Cα and 0.41–0.60 for Cγ.

### 4.2. Using multiple distinct initial conformations to improve B-factor prediction

In the above B-factor calculations, the conformational disorders of a protein crystal structure were represented by the conformational variations that resulted from 20 high-mass simulations of a protein. Specifically, each of the 20 simulations used a unique seed number for initial velocities and a common initial conformation that was taken from the protein crystal structure. These high-mass simulations were performed sequentially for (*i*) 30 ps^smt^ to set the system temperature to a desired value, (*ii*) 100 ps^smt^ to equilibrate the system at the desired temperature, and (*iii*) 25, 50, or up to 20,000 ps^smt^ to sample the atomic positional fluctuations of the protein. It was not unreasonable to suspect that the conformational heterogeneity that resulted from the heating and equilibration over a combined period of 130 ps^smt^ of the 20 high-mass simulations might be insufficient to represent the conformational disorders of the protein crystal structure.

Therefore, 20 948-ns^smt^ low-mass MD simulations using FF_12_MC were carried out for each of the four proteins to obtain protein conformations that differed from the crystallographically determined conformation. FF_12_MC was used in the low-mass simulations because it could autonomously fold Ac-(AAQAA)_3_-NH_2_ [68], chignolin [69], and CLN025 [70] in 20 NPT MD simulations 2–6 times faster than FF_14_SB, suggesting that it has a higher configurational sampling efficiency than FF_14_SB [42]. In each of the 20 948-ns^smt^ low-mass simulations for each of the four proteins, a unique seed number was used for initial velocities, and the crystallographically determined protein conformation was used as the initial conformation of the 20 low-mass simulations. For each protein, three instantaneous conformations were saved at 316-ns^smt^ intervals of each of the 20 low-mass simulations, resulting in three sets of 20 distinct instantaneous conformations saved at 316 ns^smt^, 632 ns^smt^, and 948 ns^smt^. The 20 50-ps^smt^ high-mass NPT MD simulations using FF_12_MChm described in Section 4.1 were then repeated three times under the same simulation conditions except that the initial conformations of the 20 high-mass simulations were taken from those in one of the three sets of 20 distinct instantaneous conformations that were derived from the 20 low-mass simulations.

As listed in Table 4, the differences among the B-factor RMSDs derived from using the conformations saved at 316 ns^smt^, 632 ns^smt^, and 948 ns^smt^ were marginal. Of these RMSDs, most of the RMSDs on the 50-ps^smt^ timescale are smaller than those on a shorter or longer timescale (Table 4), which is consistent with the observation described in Section 4.1. For each of the four proteins, there was a significant difference in RMSD between the B-factors derived from using the conformations of the 20 low-mass simulations and those derived from using the respective crystal structure conformation (Table 4). For BPTI and lysozyme, the RMSDs derived from the conformations of the low-mass simulations were larger than those from the respective crystal structure, and the difference (mean ± SE) was ≤2.3 ± 0.6 Å^2^ (Table 4). For GB3 and ubiquitin, the reverse was observed, and the difference (mean ± SE) was ≤2.9 ± 0.6 Å^2^ (Table 4). These results suggest that use of varied conformations from the crystal structure conformation that are sampled in 20 948-ns^smt^ low-mass simulations may slightly improve the B-factor prediction for proteins that are devoid of disulfide bonds but slightly impair the prediction for proteins with their conformations restrained by disulfide bonds.

**Table 4.** Effects of the initial high-mass simulation conformation on the root mean square deviations between experimental and calculated B-factors of GB3, BPTI, ubiquitin, and lysozyme.

| Protein<br>(Temperature) | Time<br>(ps<sup>smt</sup>) | RMSD (mean ± SE in Å<sup>2</sup>) | | | |
| --- | --- | --- | --- | --- | --- |
|  |  | IC = X-ray | IC at 316 ns<sup>smt</sup> | IC at 632 ns<sup>smt</sup> | IC at 948 ns<sup>smt</sup> |
| GB3<br>(297 K) | Cα | | | | |
| | 25 | 3.7±0.1 | 3.2±0.2 | 3.3±0.2 | 3.3±0.2 |
| | 50 | 3.1±0.2 | 3.0±0.4 | 2.9±0.2 | 3.1±0.4 |
| | 100 | 3.7±0.7 | 3.8±0.8 | 2.9±0.4 | 3.4±0.4 |
| | Cγ | | | | |
| | 25 | 9.3±0.5 | 8.8±0.6 | 8.3±0.5 | 8.8±0.6 |
| | 50 | 9.2±0.8 | 10±1 | 8.5±0.6 | 9±1 |
| | 100 | 12±2 | 13±2 | 11±1 | 12±1 |
| Ubiquitin<br>(297 K) | Cα | | | | |
| | 25 | 6.2±0.3 | 6.9±0.6 | 6.6±0.4 | 6.3±0.5 |
| | 50 | 9±1 | 7±1 | 6.1±0.8 | 6.4±0.9 |
| | 100 | 16±2 | 9±2 | 9±1 | 9±1 |
| | Cγ | | | | |
| | 25 | 7.0±0.9 | 8.2±0.5 | 7.9±0.6 | 8.1±0.6 |
| | 50 | 7.3±0.9 | 8±1 | 7±1 | 9±1 |
| | 100 | 12±1 | 9±2 | 9±2 | 10±1 |
| BPTI<br>(297 K) | Cα | | | | |
| | 25 | 5.9±0.3 | 7.1±0.2 | 6.9±0.2 | 6.4±0.3 |
| | 50 | 4.8±0.6 | 6.0±0.3 | 6.0±0.3 | 5.2±0.5 |
| | 100 | 5.2±0.8 | 4.9±0.5 | 4.7±0.8 | 4.6±0.9 |
| | Cγ | | | | |
| | 25 | 8.6±0.4 | 9.4±0.6 | 9.0±0.5 | 8.3±0.6 |
| | 50 | 8.7±0.6 | 9.2±0.9 | 9.4±0.8 | 9±1 |
| | 100 | 11±1 | 10±1 | 11±1 | 10±1 |
| Lysozyme<br>(295 K) | Cα | | | | |
| | 25 | 5.2±0.3 | 5.8±0.2 | 5.8±0.3 | 5.5±0.3 |
| | 50 | 4.2±0.4 | 5.1±0.4 | 5.2±0.7 | 4.7±0.9 |
| | 100 | 3.5±0.6 | 4.8±0.7 | 6±1 | 6±2 |
| | Cγ | | | | |
| | 25 | 7.4±0.5 | 7.9±0.7 | 7.7±0.8 | 7.9±0.7 |
| | 50 | 7.7±0.8 | 8±1 | 9±1 | 10±1 |
| | 100 | 10±1 | 10±1 | 12±2 | 14±3 |

Time: the duration of 20 distinct, independent, unrestricted, unbiased, isobaric–isothermal, and high-mass molecular dynamics simulations using FF12MChm over which the B-factors were calculated. IC: the initial conformation of a high-mass simulation that was taken either from an X-ray crystal structure or from an instantaneous conformation saved at 316 ns<sup>smt</sup>, 632 ns<sup>smt</sup>, or 948 ns<sup>smt</sup> of a low-mass molecular dynamics simulation of the respective crystal structure using FF12MC. RMSD: root mean square deviation. SE: standard error calculated from 20 distinct, independent, unrestricted, unbiased, isobaric–isothermal, and high-mass molecular dynamics simulations using FF12MChm.

### 4.3. Twenty ~50-ps^smt^ simulations might be conducive to prediction of B-factors

The present study demonstrates that the atomic positional fluctuations of a folded globular protein sampled over a timescale of ~50 ps^smt^ of 20 high-mass MD simulations can approximate the experimental B-factors better than the fluctuations sampled over a shorter or longer timescale. This observation is in agreement with a recent report showing that the experimental Cα and Cγ B-factors of GB3, BPTI, ubiquitin, and lysozyme could be best reproduced with the standard-mass NPT MD simulations with *Δt* = 0.10 fs^smt^ on the timescale of 50 ps^smt^ [42]. According to the mass scaling theory for compression or expansion of the MD simulation time described in Section 2, the standard-mass simulation with *Δt* = 0.10 fs^smt^ is equivalent to the high-mass simulation with *Δt* = 1.00 fs^smt^. Indeed, the Cα and Cγ B-factor RMSDs of all four proteins on the 50 ps^smt^ timescale in Table 2 are nearly identical to the corresponding ones in Table S14 of Ref. [42]. Further, the present finding that sampling over 50 ps^smt^ in 20 high-mass MD simulations best reproduces the experimental B-factors is consistent with the report that the internal motions are on the order of tens or hundreds of ps^smt^ [67]. It is also consistent with the report that the experimental Lipari-Szabo order parameters [71] of backbone N–H bonds of the four proteins were best reproduced with NPT MD simulations using FF_12_MC on the timescale of 50 ps^smt^ [42]. These consistent results suggest that through performing multiple picosecond high-mass NPT MD simulations one could capture the true thermal motions of folded globular proteins that are reflected in either B-factors or the Lipari-Szabo order parameters.

This study compared two simulation conditions for B-factor prediction. One used the conformational heterogeneity resulting from the heating and equilibration of a respective crystal structure over a combined period of 130 ps^smt^ of 20 high-mass MD simulations. The other used the conformational heterogeneity resulting from the heating and equilibration of multiple distinct instantaneous conformations, which were taken from 20 948-ns^smt^ low-mass MD simulations of the respective crystal structure, over a combined period of 130 ps^smt^ of 20 high-mass MD simulations. The result of this comparative study shows that sampling the atomic positional fluctuations of the simulations using multiple distinct instantaneous conformations approximates the experimental B-factors of GB3 and ubiquitin better than sampling the fluctuations of the simulations using a crystal structure conformation, and vise versa for BPTI and lysozyme. This observation correlates well with the structures of the four proteins. Unlike BPTI and lysozyme, GB3 and ubiquitin do not have any disulfide bonds to restrain their folded conformations. There is no structural difference between the solution and solid states for GB3 or ubiquitin [54,56,72,73]. However, the C14–C38 disulfide bond in BPTI flips between left- and right-handed configurations [74] in the NMR structure (PDB ID: _1_PIT) [75]. This bond is locked at the right-handed configuration in the crystal structure (PDB ID: _4_PTI) [55]. For lysozyme, its C6_4_–C80 disulfide bond adopts both configurations in the NMR structure (PDB ID: _1_E8L) [76] and the left-handed configuration in the crystal structure (PDB ID: _4_LZT) [57]. As reported recently in Ref. [42], sampling the conformation of BPTI in solution using FF_12_MC for 3.16 ns^smt^ captured both left- and right-handed configurations of C14–C38, but the left-handed configuration is absent at the crystalline state. This explains why sampling the atomic positional fluctuations over multiple distinct instantaneous conformations in solution impaired the B-factors of BPTI and lysozyme, but improved those of GB3 and ubiquitin. This also helps clarify why the B-factor RMSDs predicted using FF_12_MC progressed in time (Table 2) and underscores the necessity to confine the sampling to the timescale of ~50 ps^smt^.

In this study, the average PCCs of the predicted Cα B-factors using FF_12_MC and FF_14_SB relative to the experimental values are 0.75 and 0.74, respectively, while the individual PCCs of the predicted Cα B-factors for lysozyme using FF_12_MC and FF_14_SB are 0.79 and 0.71, respectively. To date, the best reported average PCC of the predicted Cα B-factors using a statistical method is 0.61 [46]; the best reported individual PCC of the predicted Cα B-factors of lysozyme using a single-parameter harmonic potential is 0.71 [47]. These coefficients suggest that the physics-based method that uses multiple ~50-ps^smt^ NPT MD simulations with FF_12_MC or FF_14_SB to predict Cα B-factors may be as good as if not better than the knowledge-based methods that use statistics or single-parameter harmonic potentials to predict Cα B-factors. Further, according to a survey of ~900 amino acids in four protein crystal structures with resolutions of 1.60–1.70 Å, the 95% confidence interval for the experimental B-factors derived by the refinement procedure is mean ± ~9.8 Å^2^ [8]. The present study shows that the upper limit of the RMSDs between 556 calculated Cα and Cγ B-factors (Table S1 of Supplementary Material) and the corresponding experimental B-factors of GB3, ubiquitin, BPTI, and lysozyme with resolutions of 0.95–1.80 Å is 9.6 Å^2^ (Table 2). This limit indicates that the Cα and Cγ B-factors of the four proteins predicted from 20 50-ps^smt^ high-mass simulations using FF_12_MC or FF_14_SB are accurate because these predicted B-factors are within the 95% confidence interval of the experimental B-factors.

While further studies are needed, the present work suggests that sampling the atomic positional fluctuations in 20 distinct, independent, unrestricted, unbiased, ~50-ps^smt^, and high-mass classical NPT MD simulations may be a feasible MD simulation procedure of a physics-based method to accurately predict B-factors of a folded globular protein. These high-mass simulations may use 20 distinct, initial conformations taken from the last instantaneous conformations of 20 distinct, independent, unrestricted, unbiased, 316-ns^smt^, and low-mass classical NPT MD simulations of a comparative model of the globular protein to prospectively predict main-chain and side-chain B-factors for target-structure–based drug design. These high-mass simulations may also use a common initial conformation taken from a crystal structure to retrospectively predict B-factors for insights into relative contributions of the thermal motions in time and the conformational and static lattice disorders in space to the experimental B-factors.

## Conflict of interest

The author declares no conflict of interest.

## Acknowledgments

Yuan-Ping Pang acknowledges the support of this work from the US Defense Advanced Research Projects Agency (DAAD19-01-1-0322), the US Army Medical Research Material Command (W81XWH-04-2-0001), the US Army Research Office (DAAD19-03-1-0318, W911NF-09-1-0095, and W911NF-16-1-0264), the US Department of Defense High Performance Computing Modernization Office, and the Mayo Foundation for Medical Education and Research. The contents of this article are the sole responsibility of the author and do not necessarily represent the official views of the funders. The author is most grateful to the organizers of the RapiData course at the US National Synchrotron Light Source of the Brookhaven National Laboratory, which offered him hands-on training in macromolecular X-ray diffraction measurement and inspired this work. The author also thanks two editors and an anonymous reviewer for their comments and suggestions.

